# Development of an interactive open source software application (RadaR) for infection management / antimicrobial stewardship

**DOI:** 10.1101/347534

**Authors:** Christian F. Luz, Matthijs S. Berends, Jan-Willem H. Dik, Mariëtte Lokate, Céline Pulcini, Corinna Glasner, Bhanu Sinha

**Affiliations:** University of Groningen, University Medical Center Groningen, Department of Medical Microbiology and Infection Prevention, Groningen, The Netherlands; Certe Medical Diagnostics and Advice, Groningen, The Netherlands; Université de Lorraine, APEMAC, Nancy, France; Université de Lorraine, CHRU-Nancy, Infectious Diseases Department, Nancy, France

## Abstract

**Objectives:** Analysing process and outcome measures for patients suspected of or having an infection in an entire hospital requires processing large datasets and accounting for numerous patient parameters and treatment guidelines. Rapid, reproducible and adaptable analyses usually need substantial technical expertise but can yield valuable insight for infection management and antimicrobial stewardship (AMS) teams. We describe a software application (RadaR - Rapid analysis of diagnostic and antimicrobial patterns in R) for infection management allowing user-friendly, intuitive and interactive analysis of large datasets without prior in-depth statistical or software knowledge.

**Methods and Results:** RadaR was built in R, an open source programming language, making it free to use and adaptable to different settings. Shiny, an additional open source package to implement web-application frameworks in R, was used to develop the application. RadaR was developed in the context of a 1339-bed academic tertiary referral hospital to handle data of more than 180,000 admissions.

RadaR visualizes analytical graphs and statistical summaries in an interactive manner within seconds. Users can filter large patient groups by 17 different criteria and investigate antimicrobial use, microbiological diagnostic use and results, and outcome in length of stay. Results can easily be stratified and grouped to compare individually defined patient groups. Finally, datasets of identified patients / groups can be downloaded for further analyses.

**Conclusion:** RadaR facilitates understanding and communication of trends in antimicrobial use, diagnostic use and patient outcome by linking and aggregating individual patient data in one user-friendly application. RadaR can produce aggregated data analysis while preserving patients’ features in the data to adjust and stratify results in detail. AMS teams can use RadaR to identify areas, both for diagnostic and therapeutic procedures, within their institutions that might benefit from increased support and to target their interventions.

## Introduction

In times of rising antimicrobial resistances (AMR) worldwide efforts focus on the preservation of antimicrobials as a precious non-renewable resource. Infection management in the form of antimicrobial stewardship (AMS) programmes, defined as “a coherent set of actions which promote using antimicrobials responsibly”, is an effective way to tackle this global health problem (1). Appropriate use of antimicrobials based on appropriate and timely diagnostics are important tools for successful infection management. Stewardship efforts aim at improving quality of care and patient safety as a top priority of any intervention. Thereby, the contribute to efforts in fighting AMR through optimizing the use of antimicrobials. They are focused both on individual patients (personalized medicine, consulting) and patient groups/clinical syndromes (guidelines, protocols, IT infrastructure/clinical decision support systems, etc.).

AMS setups in hospitals are often heterogeneous but audit and feedback to assess the goals are an essential part of most programmes and are included in international guidelines and reviews (2–7). Important data for AMS programmes include for example days of therapy (DOT), daily defined doses (DDD), admission dates, length of stay (LOS) and adherence to local or national treatment guidelines (1). Clinical outcomes, quality of care or consumption of hospital resources can be measured in mortality or surrogate parameters like length of stay. Widespread electronic health records (EHR) and administrative data facilitate data collection. Administrative data has also been shown to be a reliable source for assessing clinical outcomes (8).

EHR usually offer quick insights into useful data for AMS on the individual patient level. However, easy access to analyse patient groups (e.g. stratified by departments or wards, specific antimicrobials or diagnostic procedures used) is difficult to implement in daily practice. Rapid analysis of larger patient populations (e.g. spread over multiple specialties) is even more challenging although this information might be available. Yet, this is vital for meaningful analysis, including possible confounders, and pattern recognition across different populations. Even when aggregated data is available it is often not possible to trace back individual patients and analysis lack the ability to be further adjusted or stratified.

AMS teams are multidisciplinary and act beyond the borders of single specialties (9). They are usually understaffed (10), with limited IT support. They therefore need user-friendly and time-saving IT resources, without the need for profound technical expertise once the system is set up. To aggregate and link data of diagnostic compliance, antimicrobial use, guideline adherence, and clinical outcomes on the institutional level can build the basis for important insights for these teams. These could be used to identify areas within hospitals that might benefit most from supportive AMS interventions (e.g. subspecialties with lower guideline adherence, low diagnostic compliance or unusual patterns of antimicrobial use). Feedback from this data could also help physicians to better understand their patient population as a whole and administrative departments could allocate resources in a more targeted fashion.

Moreover, aggregated data and simultaneous analysis of multiple areas (e.g. antimicrobial and diagnostic use) present an extensive insight into large patient populations. This also enables comprehensive and multidisciplinary approaches of infection management combining treatment and diagnostic perspectives (1,9,11). Unfortunately, this kind of analyses still requires substantial statistical knowledge and software skills while being time consuming.

Technology, data science, and application development can bring solutions to complex data handling problems like those described above. Application development for medical and epidemiological (research) questions has found many important answers during the last years alone. Applications at hospital emergency departments (ED) in form of a software dashboard have been shown to improve efficiency and quality of care for patients requiring emergency admission to hospital. These applications are used to communicate clearly defined clinical problems like mortality ratio, number of cardiac arrests or re-admission rate to the ED (12). This approach led to a decreased length of stay and mortality at the ED. Other applications can rapidly and interactively display geographical locations of tuberculosis cases without the need of technical expertise improving the understanding of transmission and detection (13). Data driven fields like genomics are front runners in developing new innovative software applications to handle large datasets, in close collaboration with bioinformatics (14).

It is of importance to note that all of these examples have been created in an open source approach. The underlying source code can be shared, modified and freely distributed through open repositories like GitHub (https://github.com), taking open source software license obligations into account. This facilitates collaboration and easy adaptation to many different settings and IT systems and supports the use of advanced data visualization to users with minimal experience in programming and little or no budget for professional database engineers (11).

However, to the best of our knowledge, there is no such open-source approach described for applications working with antimicrobials use and diagnostic data in the field of infection management.

We followed principles of open knowledge (15) to address the need for an interactive, easy-to-use tool that allows users to investigate antimicrobial use, microbiological diagnostic use and patient outcomes at an institutional (hospital) level. We describe the development and use of a software application (RadaR - Rapid analysis of diagnostics and antimicrobial patterns in R). It enables rapid and reproducible data analysis without extensive technical expertise in a graphically appealing way while being adaptable to different settings. Analyses are based on datasets of individual patients. Therefore, also aggregated results can be stripped down and additional patient features can be investigated. With this application we aim at supporting data-driven hospital insights and decision making for actors in the field of antimicrobial stewardship.

## Methods

RadaR was developed in R, an open source programming language (16), and RStudio (https://www.rstudio.com). To build this web-browser based application we used the Shiny package for R (17). Shiny allows R users to build interactive web applications without extensive knowledge in web design and its programming languages. They can be run and hosted online for free (https://www.shinyapps.io), on local or cloud-based servers or on personal computers.

This open source approach has great advantages over traditional methods like Excel or SPSS. Hughes et al. described those in their report of an application for RNA-sequencing data analysis (14). They highlight aspects that were also fundamental for the development or RadaR. Firstly, R allows transparent, reproducible and sustainable data analysis through scripts that can easily be shared and changed. This can build the basis for collaboration and enforces the spirit of open science (also through the strong collaborative R community online). Secondly, R is open source and free to use, therefore, enabling use in resource-limited settings, too. Finally, Shiny empowers users to interact with the data making even very large datasets quickly interpretable.

The functionality of R can be easily extended by installing additional packages. Especially the dplyr and ggplot packages are to be mentioned as they are used to transform the data and to create most plots in RadaR (18,19). All packages used for the development of RadaR are listed in Table 1. RadaR is developed in an open source environment and licensed under GNU General Public License v2.0 giving options to change, modify and adapt RadaR to both personal and commercial users’ needs while requiring the need to document code changes.

**Table 1.**
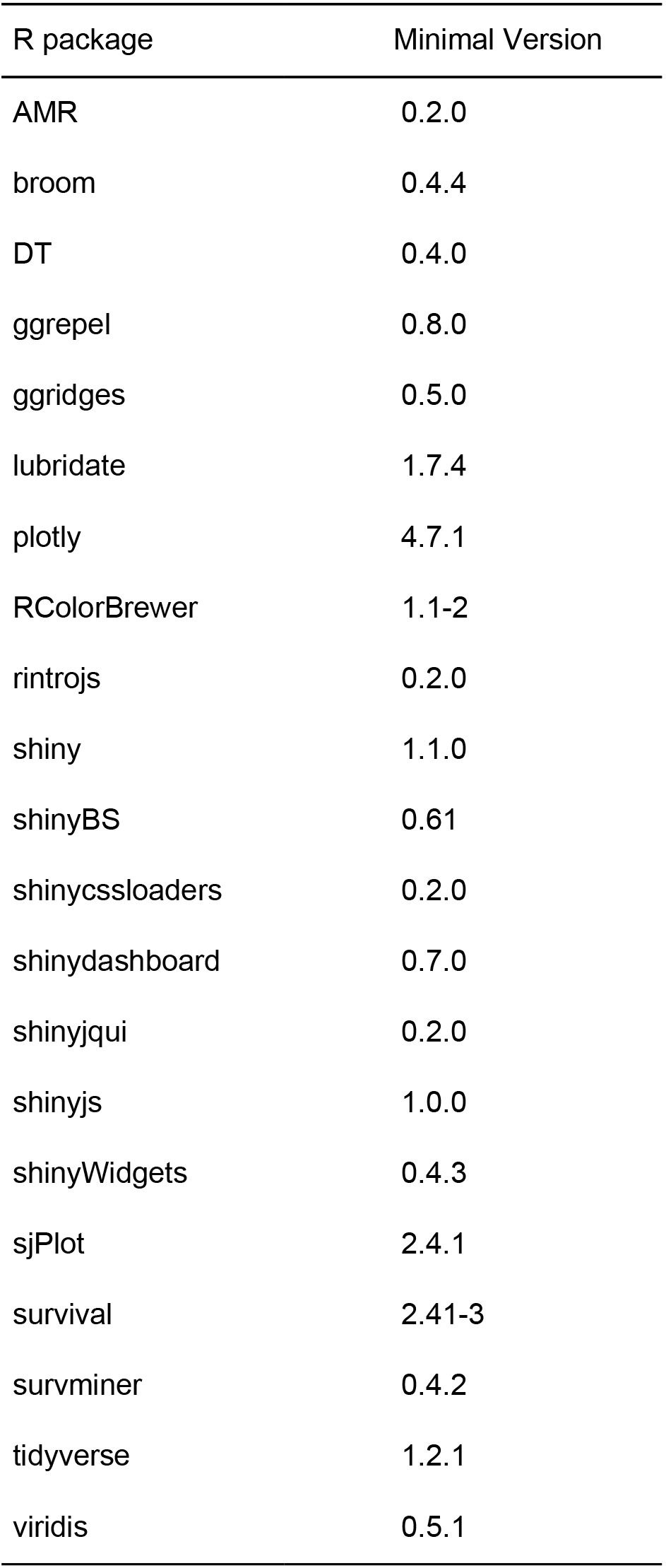
Required R packages fo RadaR

RadaR can be used to perform statistical tests for explanatory analysis. Differences in length of stay are displayed by a Kaplan-Meier curve in conjunction with a log-rank test using the survminer package (20). Users can further build a Cox regression model for length of stay and a multiple logistic regression model for the likelihood of performing microbiological diagnostic tests. The underlying calculations are done using the survival package and the base stats package respectively (16,21).

RadaR has been developed in macOS High Sierra (1.4 GHz, 4 GB RAM) and its use in Windows 7 and Linux (Ubuntu 16.04.4 LTS) was tested without any difficulties. A running example version has been deployed to shinyapps.io (https://ceefluz.shinyapps.io/radar/).

The entire source code of RadaR is freely accessible on Github (https://github.com/ceefluz/radar). The development will continue with more suggestions and improvements coming from its users and the R community. Further comments and suggestions for RadaR can be submitted at https://github.com/ceefluz/radar/issues.

## Results and discussion

We have developed RadaR, a dashboard application providing an intuitive platform for rapid analysis of large datasets containing information about patients’ admission, antimicrobial use, and results of microbiological diagnostic tests. This application can help users (i.e. professionals involved in antimicrobial stewardship programmes) to find answers to questions like: “What are the most commonly used antimicrobials at an institution/specialty/department and have they changed over time?”, “Were adequate microbiological diagnostics performed at the start of antimicrobial treatments?”, “What are the most frequent microorganisms found in different departments?”, “Can we identify priority areas within a hospital where antimicrobial or microbiological diagnostic use has the largest room for improvement?”

### Application design

RadaR is designed in form of a browser-based dashboard that most users are familiar with from typical websites and online tools (Fig. 1). The basis of RadaR’s functionality is filtering datasets and producing analytical graphs according to selection criteria defined by the user. Any calculations and data aggregation is based on single observations of individual patients. To identify and analyze groups of patients, 17 different selection criteria can be found in the sidebar (Table. 2). The output of RadaR is grouped into four panels (patient, antimicrobials, diagnostics, and outcome) that each consist of three to four output boxes displaying the results.

**Figure 1.**
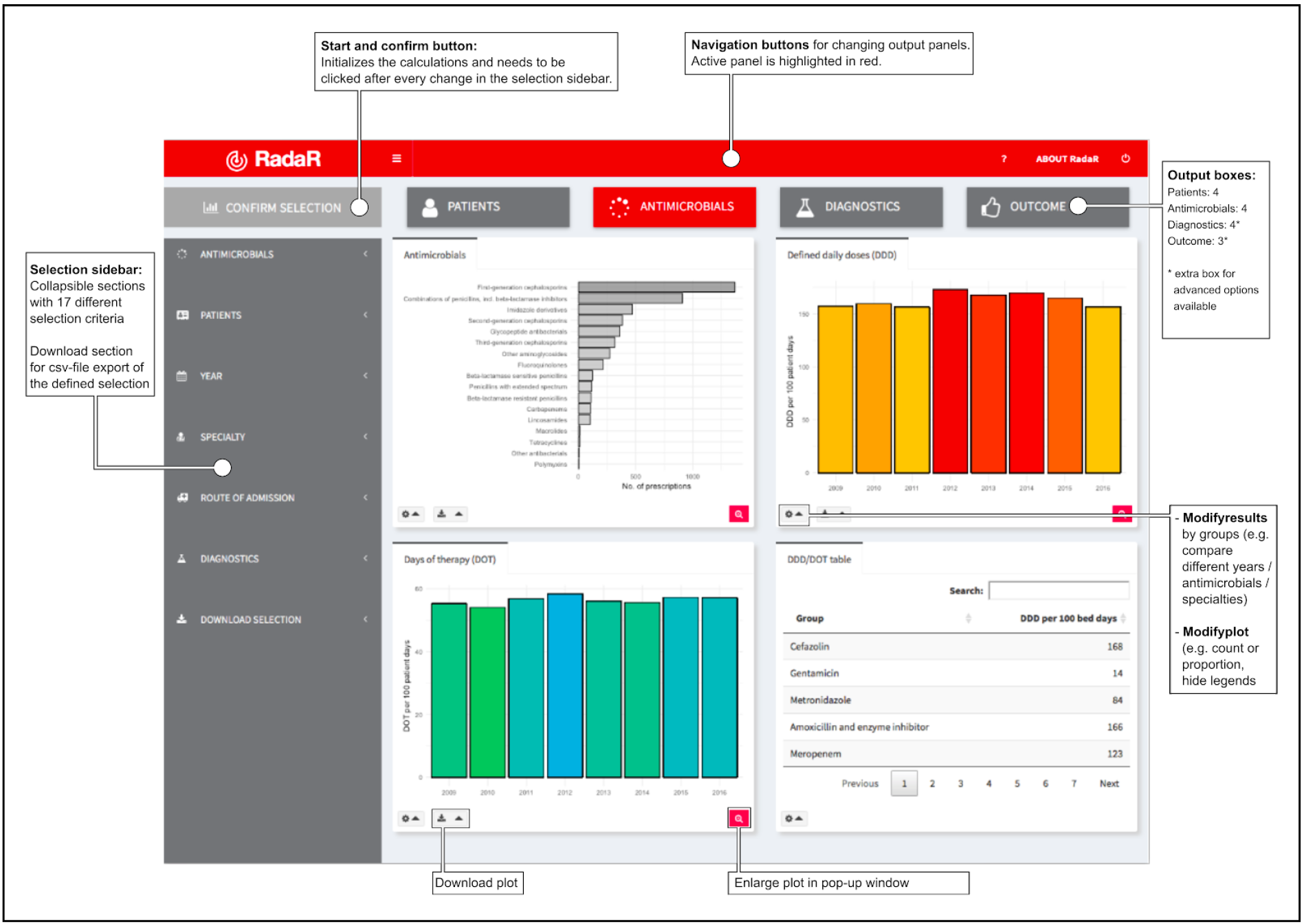
Design of RadaR in a browser-based dashboard format.

**Table 2.**
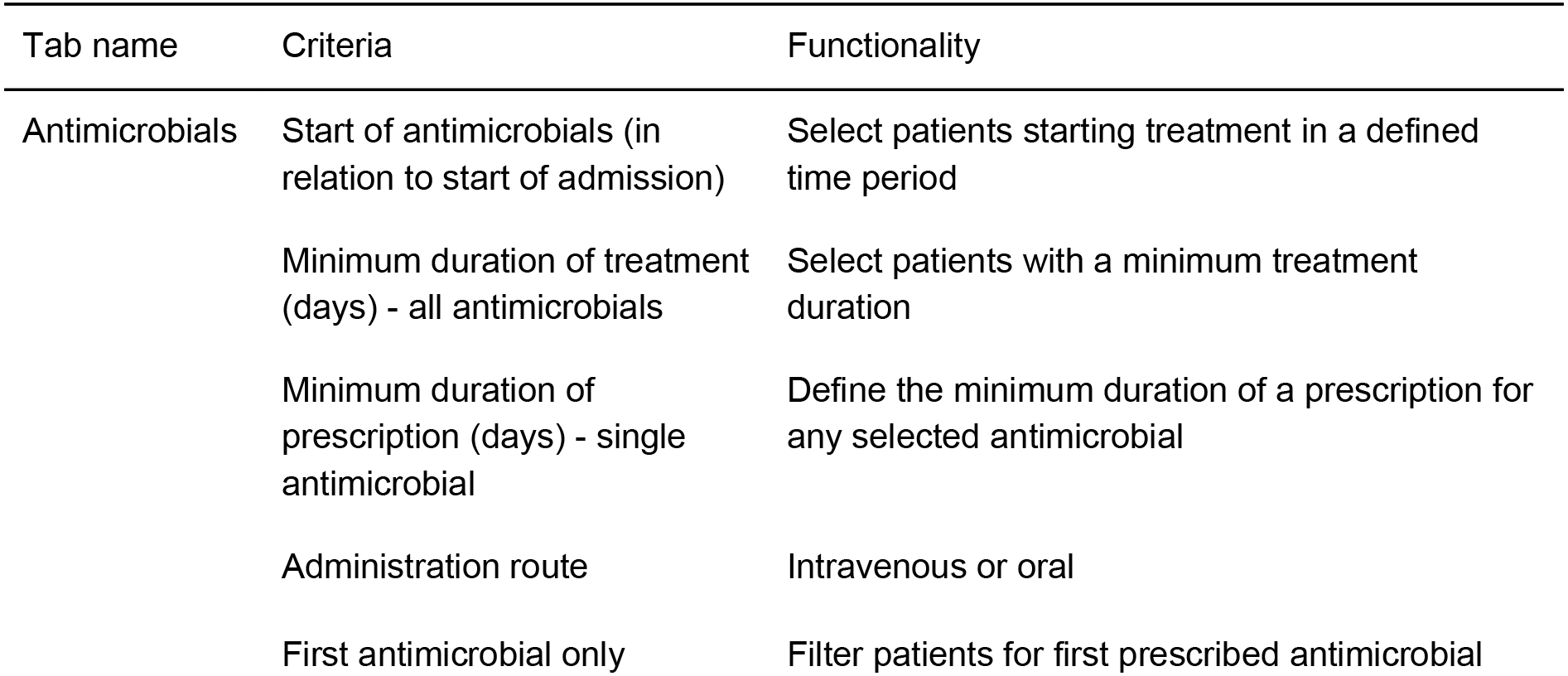
Selection criteria in sidebar.

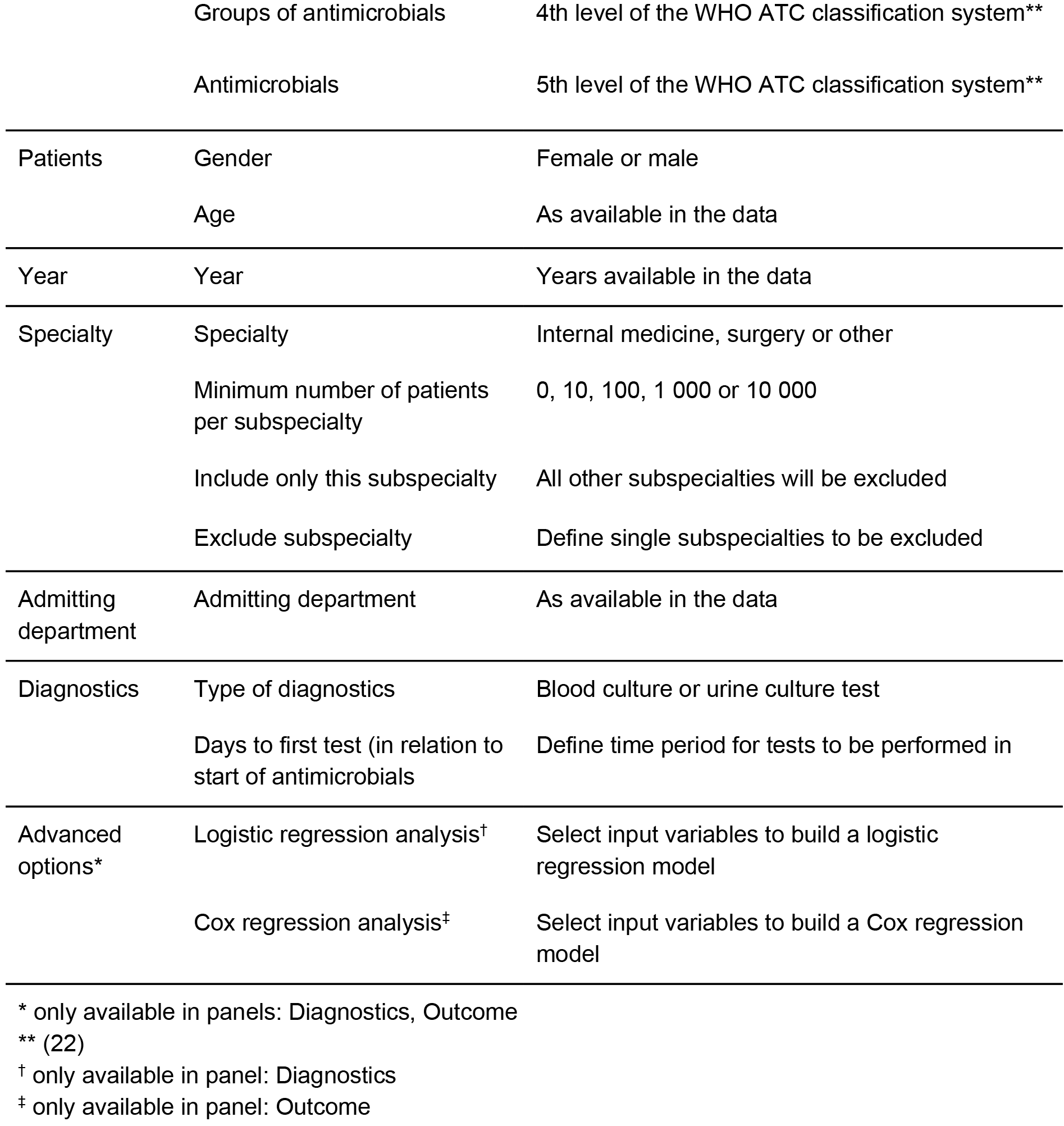

All output is based on the selection criteria defined by the user in the sidebar. By default, all available options are set to include the maximum number of patients. Each new selection and any change needs to be confirmed by clicking the confirm selection button (Fig. 1). Users can navigate between the different analysis panels through clicking the respective button. The active panel is highlighted by a change in color of the selected button.

Results are shown in bar charts, combined histogram and density plots, a bubble plot and a Kaplan-Meier curve for length of stay in hospital. Each panel further displays a table summarizing the respective data analyses. All output boxes and their content are described in Table 3. Most output boxes include modification options that can be identified by small gear icons (Fig. 1). These clickable icons allow for further specification of the generated plots and tables. Users can compare different groups (e.g. antimicrobial use by antimicrobial agent or length of stay by specialty) or modify the plots (e.g. switch from count to proportion, show or hide a histogram in density plots, show or hide the legend or highlight outliers). The combination of different options results in a total of 75 different plots that can be generated for a single selection defined in the sidebar.

**Table 3.**
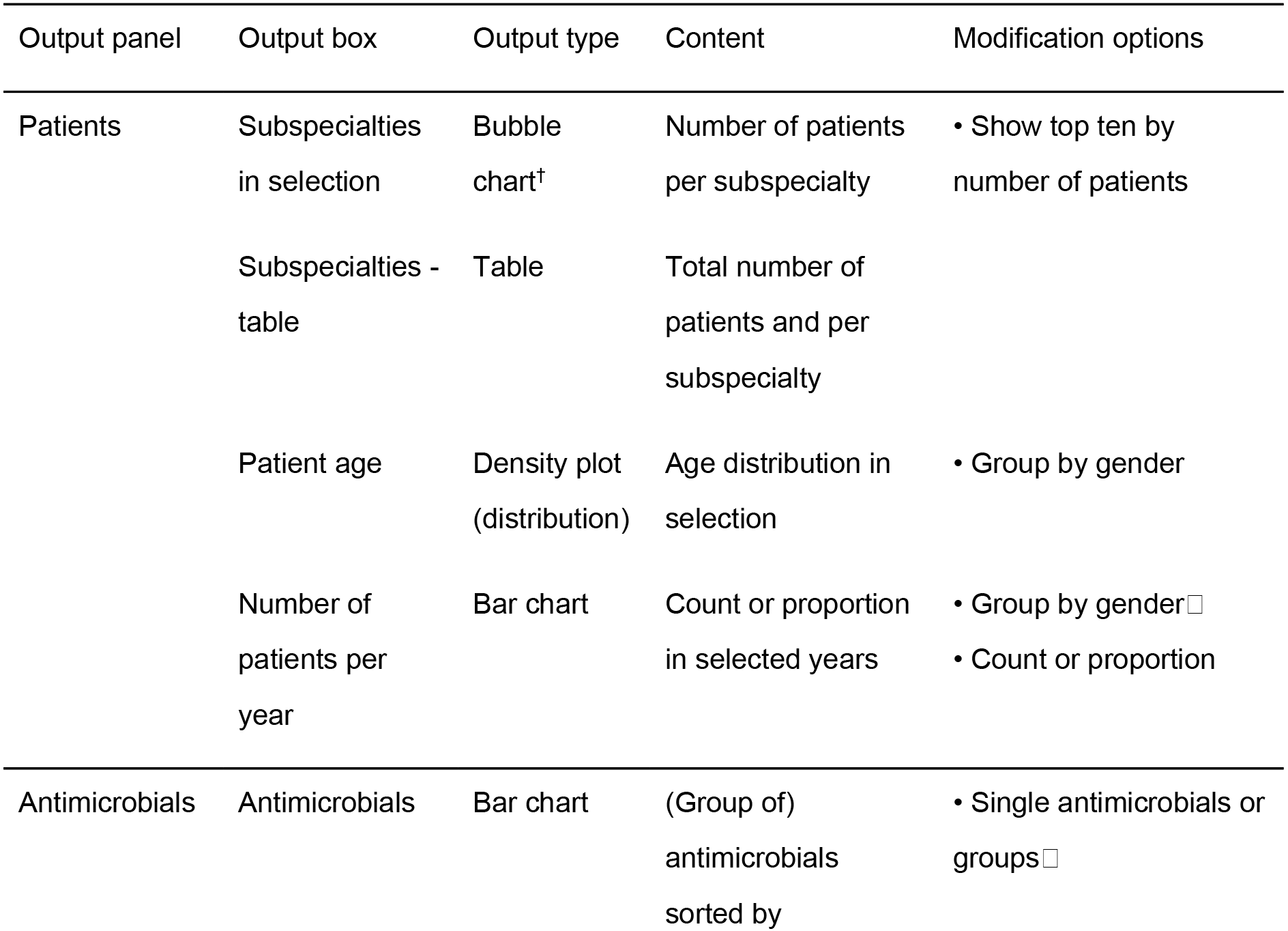
Output boxes for analysis results.

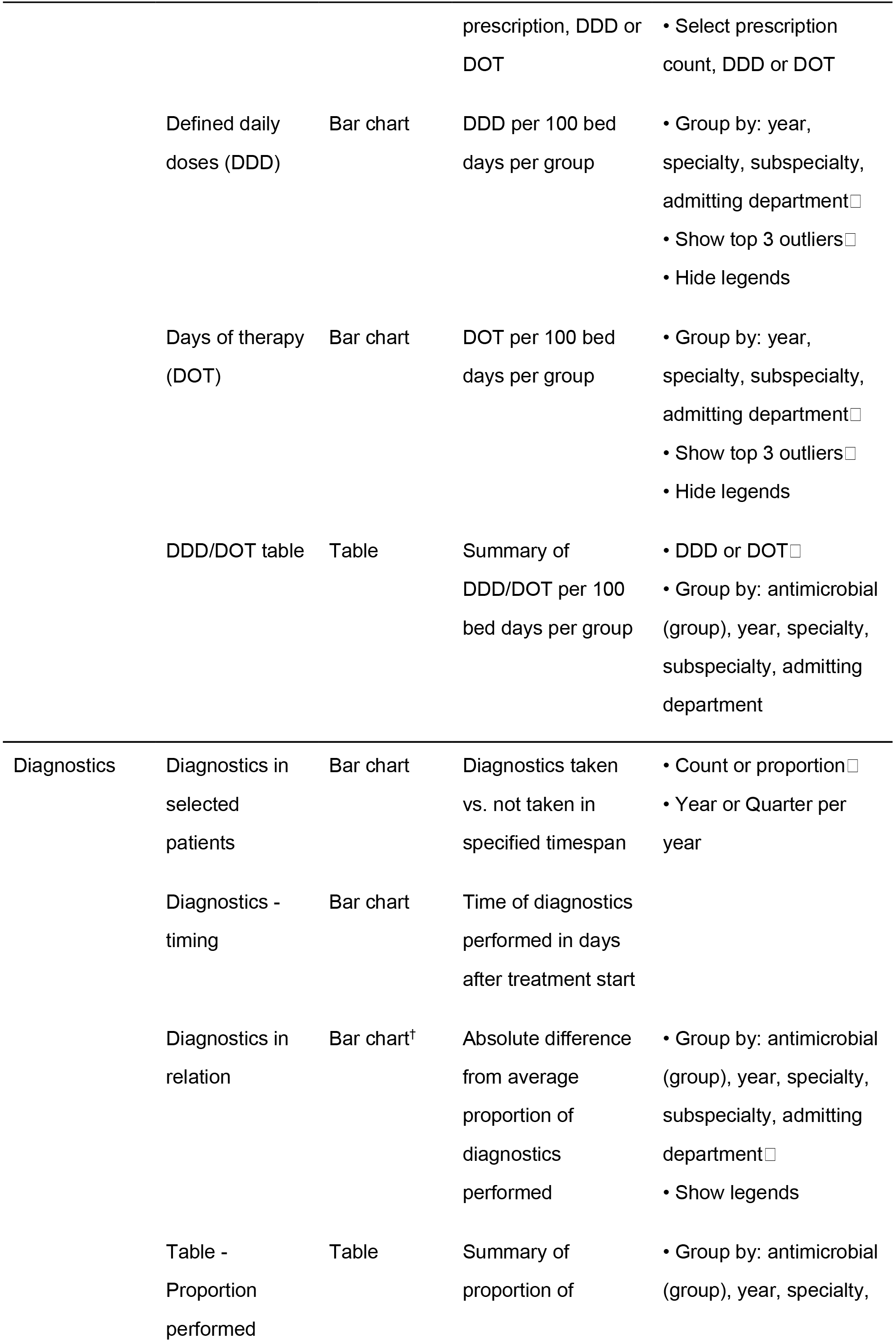

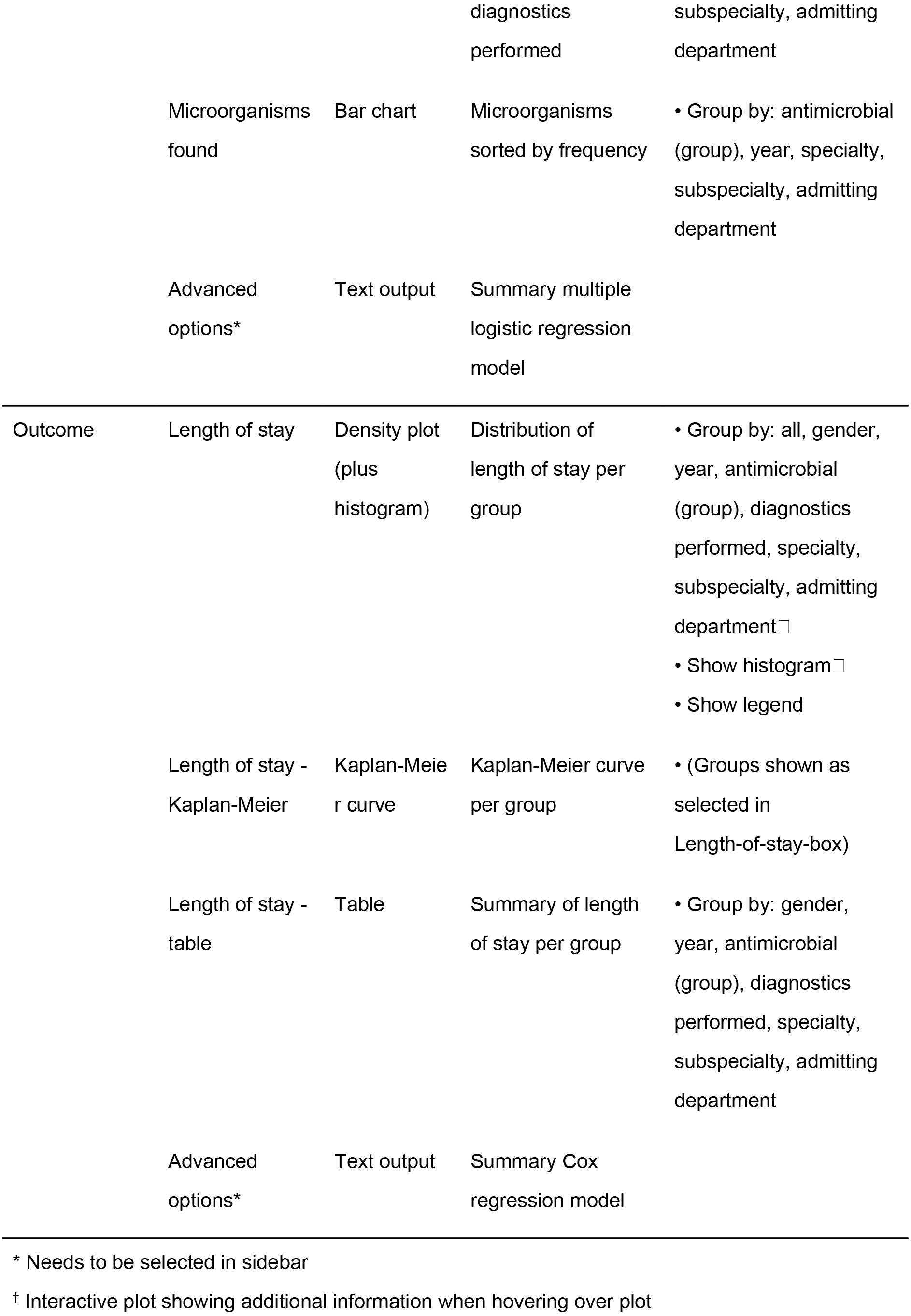

The panels for diagnostics and outcome have additional advanced options available in the sidebar. Users can use RadaR to build a logistic regression model to study the effect of different selection criteria on the likelihood of performing the selected microbiological diagnostic test in the defined time period. Similarly, the outcome panel gives the option to investigate the likelihood of discharge from hospital through a Cox regression model. The results are shown in odds and hazard ratios respectively including summary statistics to assess and compare the models. These advanced options should be used carefully and require background statistical knowledge. When implementing this application these options could be disabled depending on the local requirements.

Finally, a dataset of the user-defined selection can be downloaded from the sidebar menu in a csv-file format (comma-separated value) for further analysis (e.g. retrieving a list of patient numbers of the selected patient group)

### Development process

RadaR has been developed in close contact with infection management team at the University Medical Center Groningen, Groningen, the Netherlands, to meet the needs and requirements of this user group. Subsequently, all members of the European Society of Clinical Microbiology and Infectious Diseases Studygroup for Antimicrobial stewardship (ESGAP) were asked to evaluate and test the application through a running online example of RadaR and by filling out an online survey. The ESGAP comprises around 200 members coming from more than 30 countries worldwide. Twelve members took part in the evaluation. This yielded important information on user experiences with the application and led to further improvements that are reflected in the version we present in this report. In a next phase, RadaR will be tested in different settings of ESGAP members and other interested partners using locally available data.

### Workflow

RadaR was developed and tested with a dataset of all patients receiving systemic antimicrobials admitted to our institution, a 1339-bed academic tertiary referral hospital, within the years of 2009 to 2016 comprising 181,078 admissions with a total of 635,227 observations/rows. For simulation purposes and online user testing we have created a test dataset of 60,000 simulated patients that do not represent real patients. This dataset cannot be used for any meaningful results but allows testing of RadaR’s functionality.

A typical example workflow with RadaR comprises six steps (with examples from the test dataset):

1. Define the selection: For example, patients receiving intravenous cefuroxime for at least two days starting on the day of hospital admission from any specialty in all years in the dataset.
2. Patients panel: Identify the total number of patients and the subspecialties with the highest number of included patients (example: 188 patients selected in total with 41 patients from internal medicine and 32 from surgery). Investigate patients’ gender and age distribution.
3. Antimicrobials panel: Identify the total use of antimicrobials in DDD and DOT per 100 bed days (example: 152 and 51 respectively). Stratify the results by year and subspecialty and identify those with the highest number of DDD and DOT per 100 bed days (example: highest use by DDD and DOT in year 2011 and subspecialty pediatrics).
4. Diagnostics panel: Check if the selected microbiological diagnostic test (e.g. blood culture test) has been performed on the same day as start of cefuroxime. Investigate the proportion of performed tests over the years and which subspecialty performs best compared to others (example: surgery ICU). Check which microorganism was the most common overall and compare differences between subspecialties.
5. Outcome panel: Check for patterns of differences in LOS in the defined patient group by subspecialties or performed diagnostics (example: highest mean LOS of 12.0 days in Urology).
6. Refine the selection: Investigate a subgroup of the original selection. For example, select only the top three subspecialties by number of patients and repeat step two to five.

### Customization

RadaR uses common csv-files (comma-separated values) as input. The current state of RadaR doesn’t provide any capabilities to be immediately connected to electronic medical records. Therefore, these data still needs to be extracted manually. However, this option shall be implemented in future versions to provide the optimal use for IT staff and users.

We have documented the entire workflow of data cleaning and preparation for use with RadaR in an online accessible Rmarkdown-file (https://github.com/ceefluz/radar/blob/master/RadaR_data_preparation.Rmd). These steps are based on data from our institution and can be adjusted to fit the needs of different settings. The data should be structured in a dataset format where each variable is one column and each observation is one row. This follows the concept of tidy data by Hadley Wickham (23). Table 4 displays the final set of variables underlying RadaR’s functionality.

**Table 4.**
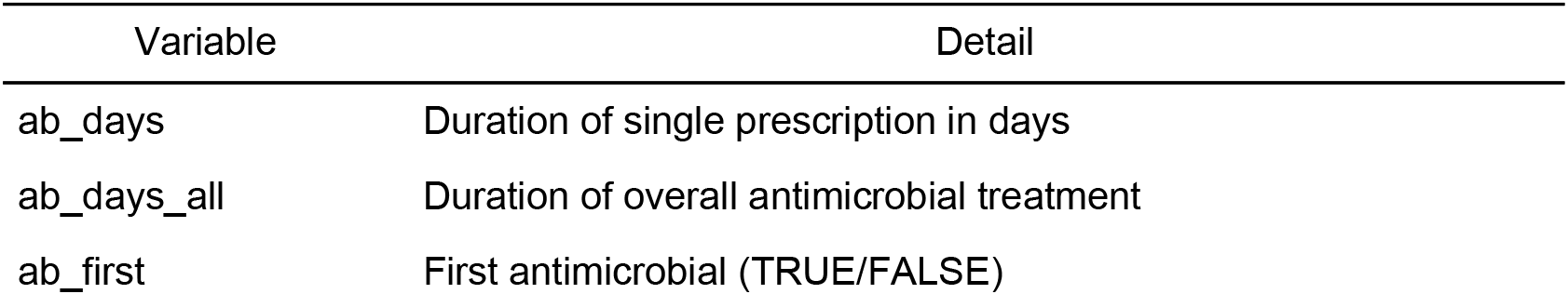
Input variables for RadaR

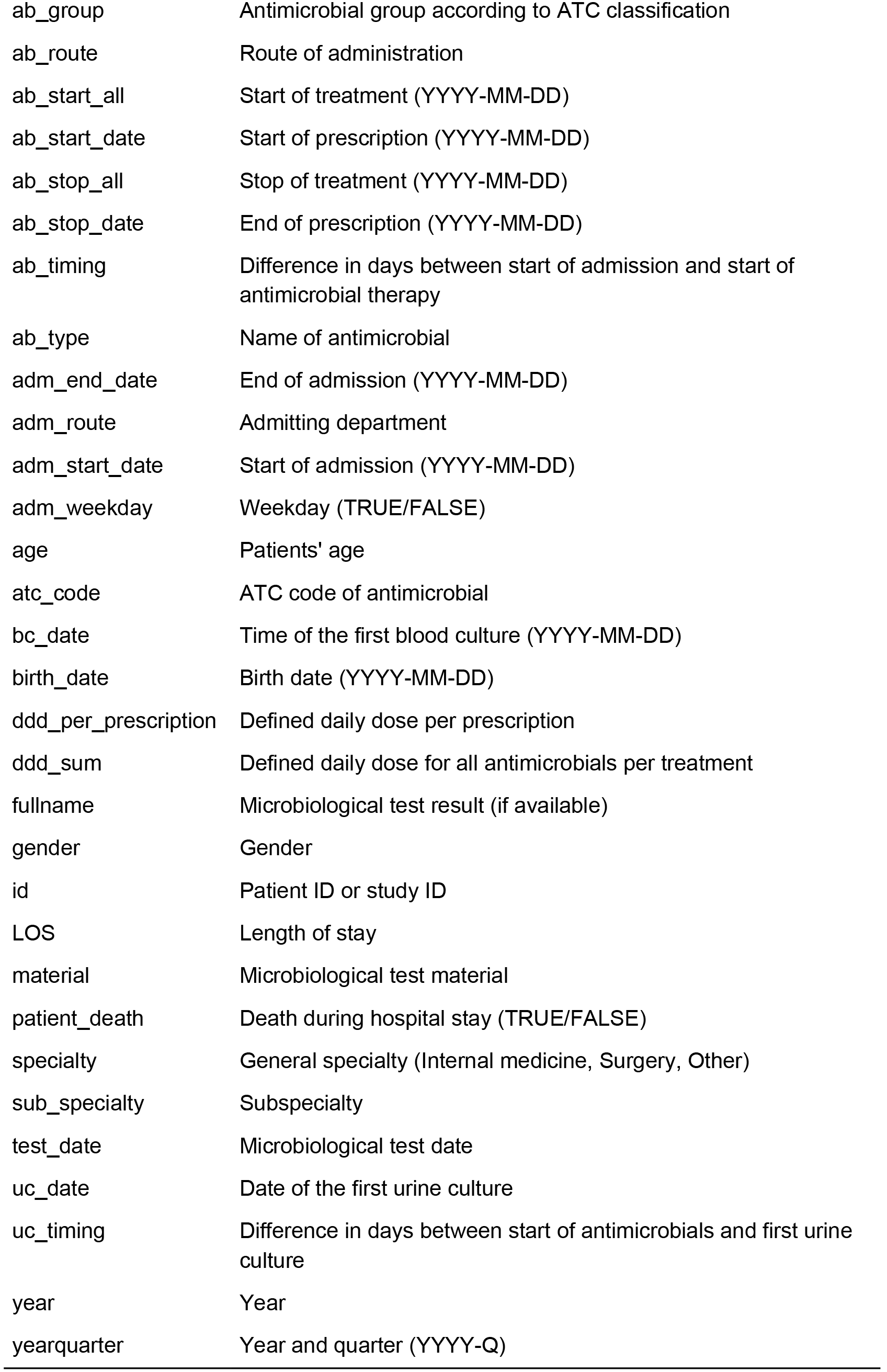

For setting up RadaR in a new environment after data preparation, users only need to perform the following four steps:

1. Downloading R (https://cran.r-proiect.org/) and RStudio (https://www.rstudio.com/products/rstudio/download/), which are free to use and open source software.
2. Download or copy and paste RadaR’s source code (https://github.com/ceefluz/radar) into three files in RStudio: global.R, server.R, and ui.R.
3. In global.R manually edit the path for the prepared data to be imported into RadaR. Only a single line of code needs to be edited.
4. Run the application in RStudio with the calling the function runApp() in the console or by clicking the green run app button. This will download and install the required R packages needed for the application if they have not been installed previously and create the final dataset for analysis. The RadaR interface will open in the RStudio viewer pane or in a new window of the standard browser of the user’s operating system.

RadaR’s appearance has been customized with using a CSS (Cascading Style Sheets) script (https://github.com/ceefluz/radar/blob/master/www/radar_style.css) that is loaded into the application upon its start. This script needs to be saved into a subdirectory of the directory of the three main files (global.R, server.R, ui.R) called “www”. We recommend RStudio’s project function to create a single project for RadaR and to store all information in this project directory. Users with experience in using CSS can fully alter RadaR’s design by changing the underlying CSS script.

### Ethical considerations

RadaR has been developed using data of patients admitted to the University Medical Center Groningen, Groningen, The Netherlands. Data was collected retrospectively and permission was granted by the ethical committee. RadaR can be used locally in protected environments. When deploying it as an online application measures have to be taken to guarantee data protection depending on national regulations.

## Conclusion

We have developed a web-browser based application for rapid analysis of diagnostic and antimicrobial patterns that can support AMS teams to tailor their interventions. It has been designed to enhance communication of relevant findings while being easily accessible also for users without prior extensive software or statistical skills. In experience this system can be adapted to new settings within one day when the required data is available. RadaR has been developed in R, an open source programming language, making it free to use, share or modify according to different needs in different settings. Next steps will involve testing in multiple settings and forming a user and research group to continue and expand the use of open source technology and open science principles in infection management. This can lead to more transparency enhancing optimal quality of care and patient safety which is crucial in the light of new data-driven developments of using EHR (24).

## Acknowledgements

We thank the ESGAP executive committee for supporting the evaluation of RadaR in the ESGAP study group and all its members for their valuable input, suggestions, and comments. Furthermore, we thank Igor van der Weide, Jan Arends and Prashant Nannan Panday for their great support in obtaining required data at our institution that built the basis for the development of RadaR. And, not to forget, the online R community that made this work possible and we are happy to contribute to the community by presenting our work on RadaR.

RadaR was developed as part of a project funded by the Marie Skłodowska-Curie Actions (Grant Agreement number: 713660 - PRONKJEWAIL - H2020-MSCA-COFUND-2015).

